# Single-cell transcriptomic analyses unveil exercise-induced changes in murine skin vascular endothelial cells

**DOI:** 10.1101/2025.05.13.653788

**Authors:** Pegah Hejazi, Pauline Mury, Éric Thorin, Guillaume Lettre

## Abstract

Physical activity (PA) is a fundamental aspect of preventive medicine, offering profound benefits for cardiovascular health and overall well-being. Despite its widespread benefits, the molecular mechanisms underlying PA-induced improvements in microvascular functions remain poorly understood. The skin microvasculature is uniquely affected by exercise-induced shear stress, especially during thermoregulation. Using single-cell RNA sequencing, we investigated how voluntary exercise influences the transcription profile of endothelial cells in the skin microvasculature of mice. We assigned 20 mice to either a sedentary group or a 1-month voluntary exercise program involving running wheels. Post-intervention, we collected skin biopsies from twelve mice for transcriptomic analyses. The differential expression analysis showed a significant increase in the expression of the *Zbtb16* gene in vascular endothelial cells (vECs). Additionally, Gene Set Enrichment Analysis (GSEA) with nominally differentially expressed genes in vECs highlighted the suppression of pathways related to oxidative stress, cell proliferation and metabolism in the exercise group. This suggests an exercise-triggered transition towards improved metabolic efficiency and enhanced homeostasis in vECs. These results begin to reveal how exercise induces molecular changes in vECs of the skin microvasculature, highlighting the role of PA in influencing endothelial function.

## INTRODUCTION

Regular physical activity (PA) significantly reduces mortality rates, decreases hospitalization frequency and enhances the quality of life for all individuals, including those with cardiovascular diseases (1). Longitudinal research has shown that maintaining or adopting an active lifestyle significantly lowers the risk of all-cause and cardiovascular disease mortality in patients with coronary heart disease, emphasizing the importance of PA trajectories in long-term health outcomes (2). Despite the known health benefits, the molecular mechanisms that explain the protective effects of PA are still not clearly understood, especially the differences among inter-individual responses.

Recent studies have shown the positive impact of PA on microvascular functions. A 8-week aerobic training program improved microcirculation reactivity and endothelial function in the skin of patients with ischemic heart disease (3). An elevated risk of coronary heart disease is associated with impaired endothelium-dependent vasodilation and reduced capillary recruitment in the skin, suggesting that skin microvascular function serves as an effective model for exploring the relationship between cardiovascular risk factors, microvascular health and PA (4).

High-throughput single-cell RNA-sequencing (scRNAseq) methods provide unique insights at cellular resolution, revealing cellular heterogeneity, differential gene expression responses, and cellular functions within highly organized tissues. These studies have already uncovered vascular cell heterogeneity in the human skin, showing that vascular endothelial cells (vECs) are more active in intercellular crosstalk rather than merely serving as passive components of the vascular lining (5,6). vECs are particularly relevant to monitor PA-induced changes in transcriptomic profiles of the skin microvasculature. It is known that elevated shear stress is the main signal that triggers endothelial adaptation to exercise, and these changes are inclusive not only to the muscle microvascular system but also to other organ such as the skin (7). During exercise, the blood flow of the working muscles increases and is primarily directed to the skin to enable thermoregulation (8). Different studies also support the notion of exercise-induced changes in vascular responsiveness of the skin (9). For instance, Wang and collaborators detected improved vascular endothelial responses, specifically enhanced endothelium-dependent dilation in the skin vasculature, following an 8-week PA training program in healthy men, with these effects reversing to the pretraining state upon detraining (10).

These findings prompted us to examine the transcriptomic profile of the murine skin microvasculature after exercise to better understand the cellular mechanisms underlying microvascular function. We focused on the cellular heterogeneity of the skin microvascular network and how exercise influences endothelial function, aiming to uncover pathways through which PA promotes vascular health. This research may provide insights into the mechanisms that modulate vascular health through PA and contribute to the development of targeted therapeutic strategies for improving the microvascular state.

## METHODS

### Animals

Twenty wildtype C57Bl/6 mice were included in this project. Among them, 10 were male and 10 were female, all were aged 5 months. All animal experiments were performed in accordance with the *Guide for the Care and Use of Experimental Animals of the Canadian Council on Animal Care* and the *Guide for the Care and Use of Laboratory Animals of the US National Institutes of Health* (NIH Publication No. 85-23, revised 1996). Experiments were approved by the Montreal Heart Institute Ethics Committee (ET No. 2019-42-01). Mice were kept under standard conditions (24°C; 12-h:12-h light/dark cycle) and were fed *ad libitum* with regular chow (2019S; Harlan Laboratories, Madison, WI, US).

### Exercise protocol

Ten mice (5 male and 5 female) were randomly assigned to the “Physical Activity” (PA) group and were exposed to 1-month of voluntary exercise. To this end, mice were kept individually in cages instrumented with a running wheel (Lafayette Instrument Company, Lafayette, IN). Each running wheel was equipped with a counter to track the running activity of each individual mouse. The remaining 10 mice were kept in standard cages (i.e. without running wheels) as sedentary controls (SED). Mice were weighted at the beginning and end of the PA month. Mice were sacrificed at 6-month after anesthesia with a 1:1 mixture of xylazine (Bayer Inc, Toronto, ON, Canada) and ketamine hydrochloride (Bioniche, Belleville, ON, Canada) at the same time of the day (morning).

### Free fatty acid plasmatic level assay

In order to validate our 1-month voluntary exercise program, we assessed free fatty acid plasma levels using the Free Fatty Acid Quantification Colorimetric/Fluorometric Kit (BioVision, Milpitas, CA).

### Vascular reactivity

Two 2-mm long segments of freshly collected mesenteric arteries were mounted in a wire myograph filled with 10 mL of physiological salt solution, as previously described (11). We recorded isometric changes in tension: arterial segments were pre-constricted with phenylephrine (PE; 10 to 30µM); at the plateau of the constriction, segments were relaxed by cumulative addition of increasing concentrations (from 1nM to 10µM) of acetylcholine (ACh) to assess endothelium-dependent relaxation. At the end of the experiment, the segment was maximally constricted with 127mM KCl-physiological solution (NaCl replaced by KCl to induce maximal depolarization of the vascular smooth muscle cells) to calculate the percent of constriction induced by phenylephrine. The concentration of ACh inducing 50% of relaxation (ACh-EC_50_), indicative of the vascular sensitivity to ACh, as well as the maximal relaxation (E_max_), were calculated to characterize endothelial function.

### Skin biopsy and scRNAseq library preparation and sequencing

Upon sacrifice, a piece of skin from the leg was harvested and immediately processed as detailed below. Two pieces of 1cm^2^ of skin were harvested per mouse and were dissociated using the Whole Skin Dissociation Kit (Miltenyi Biotec, Germany) with some variations regarding the manufacturer’ instructions. Skin tissue was incubated at 37°C in a water bath for 1h30 in the enzyme mix. Mechanical dissociation was then processed using the ThermoMixer (Eppendorf) for 1-hour at 37°C and 400 rpm. Following completion of the dissociation, the homogenate was filtered through a 70-µm cell strainer into a 50-mL centrifuge tube. The cell suspension was then centrifuged at 600g for 10 min at 4°C. Once complete, the supernatant was removed and the pellet was incubated in 500 µL of Red Blood Cell Lysis Buffer for 1min30 (Roche). Cells were again centrifuged at 600g for 10 min at 4°C and then resuspended in 500 µL of PBS + 0.04% BSA. Cell count and viability were estimated using the Countess II FL cell counter (Thermo Fisher).

Endothelial cells were then enriched using the CD31 MicroBeads mouse Kit (Miltenyi Biotec, Germany), following the manufacturer’ instructions. Briefly, cells were incubated for 30-min with 10µL of beads. After a wash step, cells were loaded onto an MS column on the OctoMACS Separator. The enriched fraction of CD31+ cells were re-counted using the Countess II FL cell counter (Thermo Fisher) and were immediately loaded onto a 10X Chip and processed on the 10X Chromium Controller. Samples were sequenced at Genome Quebec’s Centre d’Expertise et de Services on an Illumina NovaSeq sequencer with a PE100 protocol.

### Data processing and quality-control

FASTQ files were aligned to Cellranger’s mm10-2020-A reference using the count function of cellranger-7.0.1 for each sample (https://www.10xgenomics.com/support/software/cell-ranger/latest/release-notes/cr-reference-release-notes). To remove the ambient RNA contamination, we used the SoupX tool (12). For downstream analyses, we used Seurat v4 (13). We removed low-quality cells using the following thresholds: >200 and <5000 detected genes, mitochondrial reads <20, ≥500 and <40,000 sequence reads, RNA complexity >0.8. To normalize and regress out mitochondrial read percentage, we used SCTransform (14). We integrated the data using Harmony on 30 principal components (15). To annotate cell types, we obtained the skin canonical markers from two different databases: *Tabula muris* (16) and immunesinglecell skin atlas (17). We plotted the expression levels of key marker genes obtained from both databases onto the UMAP representation of our dataset to assign cel-type to each cluster. We identified doublets using scDblFinder (18). We removed cells for which the scDblFinder score was >0.5. We sub-clustered each main cell-type and removed subclusters showing doublet enrichment.

### DiNerential gene expression analysis

To identify differentially expressed genes (DEGs) across different condition groups for each cell-type, we conducted a pseudobulk analysis. This approach involves aggregating gene expression counts at the sample level within each cell-type to create bulk-like profiles for each condition. We utilized Seurat’s AggregateExpression() function to perform this aggregation, effectively summarizing the expression data for each cell-type across all samples. We used DESeq2 to perform differential expression analysis (19). Specifically, we used the likelihood ratio test (LRT) provided by DESeq2, incorporating the condition of the mice (with or without PA) as the primary predictor variable while controlling for sex as a covariate. This allowed us to identify genes that are differentially expressed in the group of mice subjected to PA compared to those that were not. Genes with a false discovery rate (FDR) <5% were significantly associated with PA.

### Gene Set Enrichment Analysis (GSEA)

To gain insights into the biological pathways and processes associated with the DEGs in vECs, we performed GSEA (20). To employ GSEA, we used the complete output from our differential gene expression analysis performed with DESeq2. This approach included both statistically significant and non-significant genes ranked based on a combined metric of their P-values and log_2_ fold change, allowing us to account for both the statistical significance and the direction of expression changes. We employed the fgseaSimple() function from the fgsea package (21) to conduct the GSEA, leveraging the hallmark gene sets from the Molecular Signatures Database (MSigDB) collection specific to mouse genes. We used the Spearman’s correlation test to identify genes that are nominally differentially expressed in vECS and correlated with the expression of *Zbtb16* (adjusted P-value <0.05).

## RESULTS

### Voluntary exercise decreased body weight and free fatty acid levels

We recorded the voluntary running mileage of all mice with an automated wheel-attached counter. We calculated the total distance run (30-days). The average run distance was 164.4 ± 24.25 km, denoting a large variability between individuals, due to the chosen mode of exercise (voluntary). To validate the efficacy of the intervention, we assessed two different parameters: (1) evolution of body weight before and after intervention and (2) plasma levels of free fatty acid (FFA) (**Fig. 1**). Evolution of body weight from mice in the PA group were significantly lower than mice in the SED group (PA = −0.06 ± 0.52 gr *vs.* SED = 1.46 ± 0.39 gr, two-tailed *t*-test P-value = 0.031) (**Fig. 1A**). Similarly, plasma levels of FFA were significantly lower in PA mice than in SED mice (PA = 0.56 ± 0.019 µM *vs.* SED = 0.62 ± 0.021 µM, P-value = 0.033) (**Fig.1B**). These two results validate our model of exercise.

**Figure 1.**
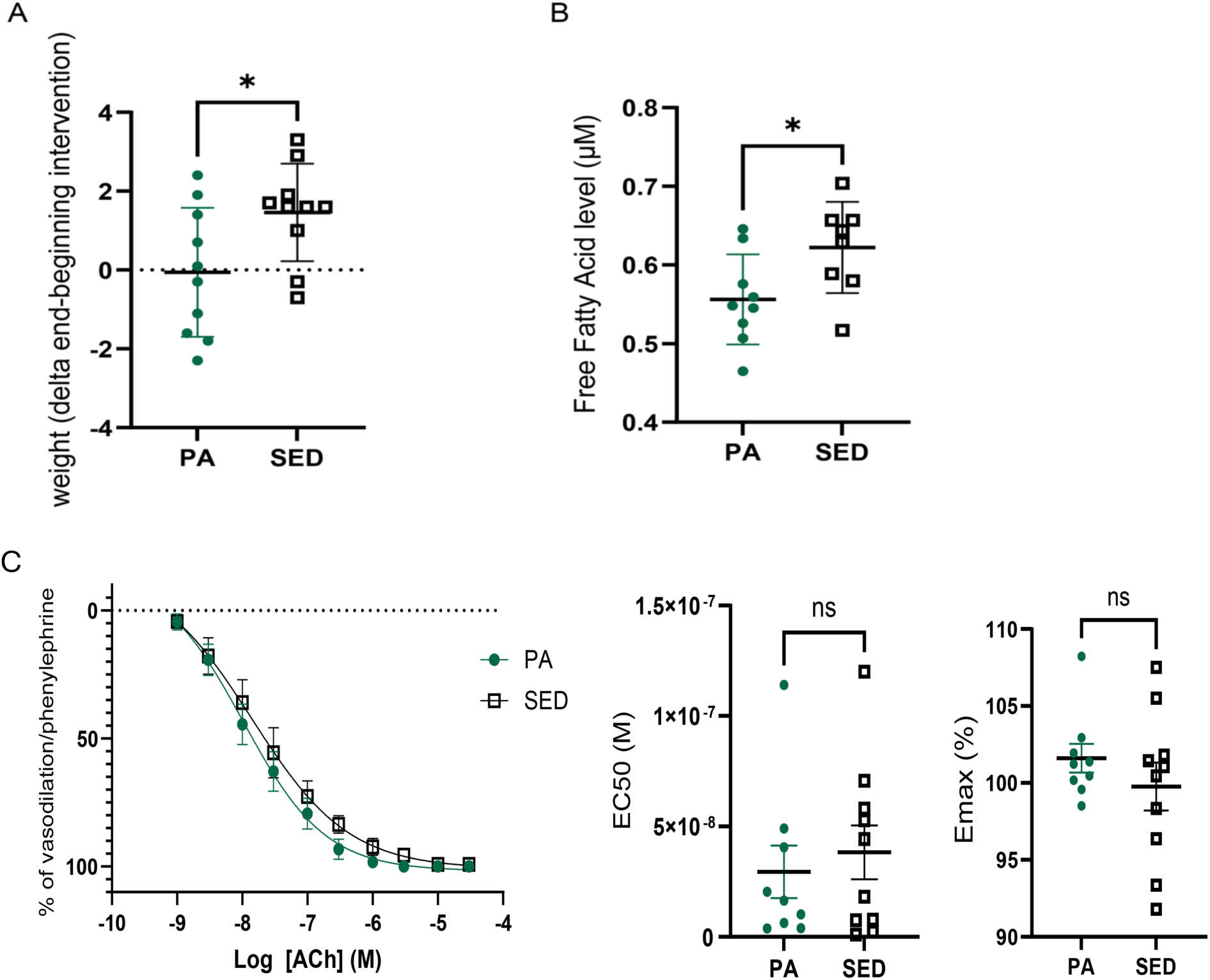
Impact of physical activity (PA) on body weight and free fatty acid levels. (**A**) Evolution of body weight from mice in the PA group that is significantly lower than mice in the sedentary group (SED)(PA = −0.06 ± 0.52 gr *vs.* SED = 1.46 ± 0.39 gr, two-tailed *t*-test P-value = 0.031). (**B**) Plasma levels of free fatty acid (FFA) that are significantly lower in PA mice than SED mice (PA = 0.56 ± 0.019 µM *vs.* SED = 0.62 ± 0.021 µM, P-value = 0.033). (**C**) Voluntary PA did not significantly improve the endothelial sensitivity to acetylcholine (ACh) (PA: 16.4 nM [5.13 – 44.8 nM] *vs.* SED: 31.2 nM [6.24 – 61.2 nM], P-value = 0.66) nor the maximal relaxation (PA: 101.6 ± 2.81% *vs.* SED: 99.8 ± 4.94 %, P-value = 0.34) after 1 month of voluntary PA. ns, non-significant.

### Vascular reactivity

We measured two indexes of vascular reactivity to assess endothelial function: ACh-EC_50_ for endothelial sensitivity to acetylcholine (ACh) and E_max_ for the maximal relaxation (**Fig.1C**). We performed the vascular reactivity analysis in each of the 20 mice, and then compared the effect of the exercise treatment. Voluntary exercise did not improve neither endothelial sensitivity (EC_50_) to ACh (PA: 16.4 nM [5.13 – 44.8 nM] *vs.* SED: 31.2 nM [6.24 – 61.2 nM], P-value = 0.66) nor maximal relaxation (E_max_) (PA: 101.6 ± 2.81% *vs.* SED: 99.8 ± 4.94 %, P-value = 0.34). This finding suggests that 1-month of voluntary exercise is not sufficiently potent to induce significant changes in arterial endothelial function that can be measured in 20 healthy 6-month-old mice.

### Identification and characterization of cell populations in murine skin

While we could not detect a physiological change in endothelial function after 1 month of exercise, we wondered whether such treatment could modify gene expression programs. To address this question, we performed scRNAseq on a subset of the same mice. Unsupervised Seurat-based clustering of 141,226 cells (48% of which are keratinocytes) from the skin of 12 mice revealed nine distinct cell-types. After quality-control steps and the removal of keratinocytes and basal cells to focus on cell-types of the microvascular environment, we obtained a single-cell dataset with 66,112 cells (**Fig. 2A**). This dataset included vECs (5.4%), smooth muscle cells (SMCs, 3.9%), fibroblasts (FBs, 31.8%), lymphatic endothelial cells (lECs, 21.0%), monocytes (31.2%), mast cells (0.6%), Langerhans cells (2.0%) and lymphocytes (4.2%)(**Fig. 2B**).

**Figure 2.**
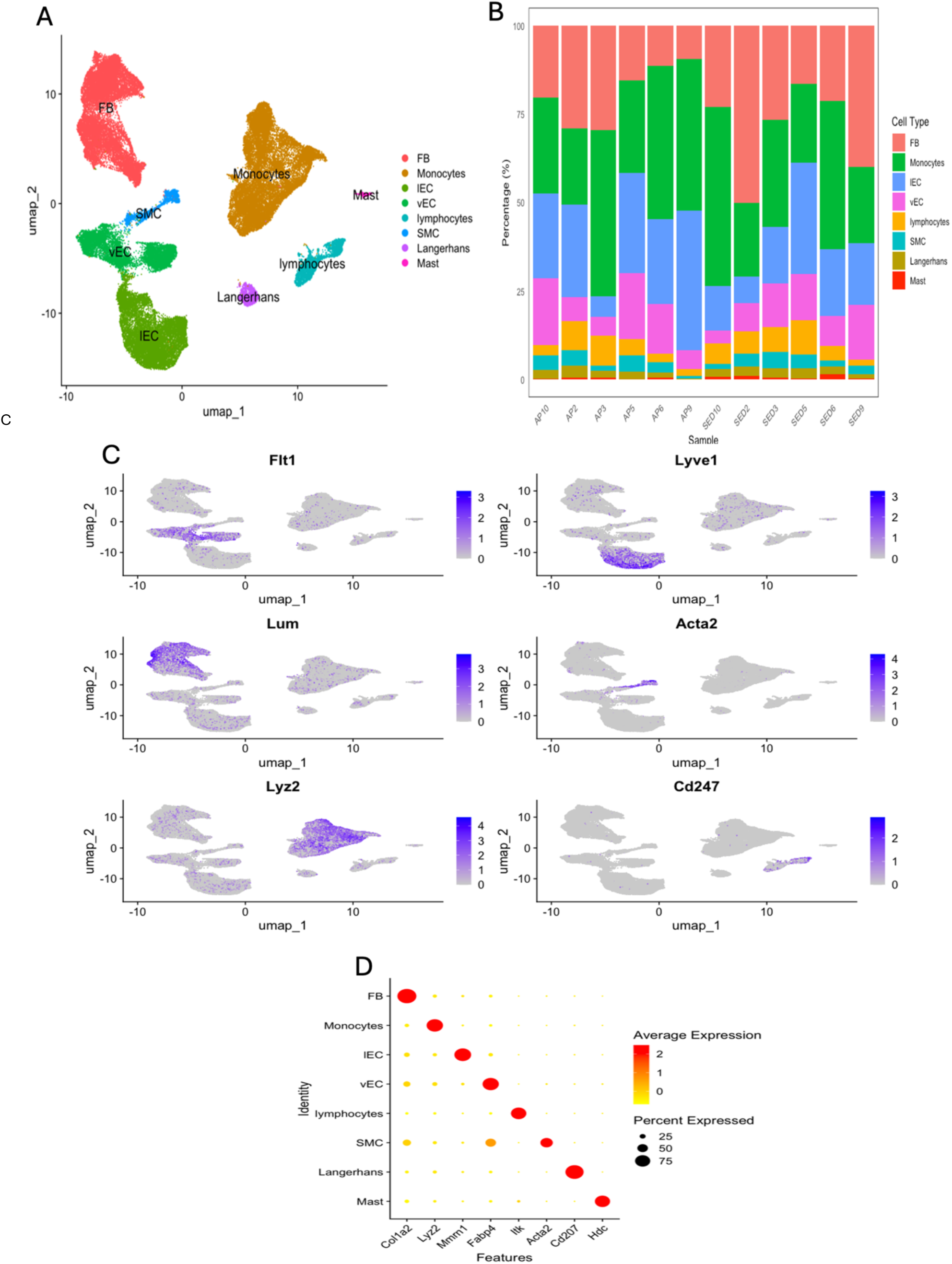
Single-cell RNA-sequencing analysis of the mouse skin microvasculature. **(A**) UMAP visualization of the dataset post-removal of keratinocytes and basal cells. The unbiased clustering of 66,112 cells displays a variety of vascular and immune cell-types. Key cell-types identified in the dataset include vascular endothelial cells (vECs), smooth muscle cells (SMCs), fibroblasts (FBs), lymphatic endothelial cells (lECs), monocytes, mast cells, Langerhans, and lymphocytes. (**B**) Stacked bar plot illustrating the proportion of di_erent cell-types across samples. Each bar represents a sample, labelled by sample IDs, with colours indicating di_erent cell types: FB (salmon pink), Langerhans (olive), IEC (light blue), monocytes (green), mast cells (red), SMC (turquoise), lymphocyte (orange), and vEC (pink). The y-axis shows the percentage of cells, highlighting the distribution and relative abundance of each cell-type within the samples. (**C**) The figure presents feature plots for key marker genes specific to each cell-type used to confirm cluster annotations. The expression of these genes was visualized across the UMAP. Each plot shows the spatial distribution and expression intensity of the marker genes, enabling the validation of cell-type identities within the clusters. Some of the key marker genes include: *Flt1*: marker for vECs, *Lyve1*: marker for lECs, *Acta2*: marker for SMCs, *Lum*: marker for FBs, *Cd247*: marker for lymphocytes, *Lyz2*: marker for monocytes. (**D**) Dot plot illustrating the primary marker gene for each cluster by average and percentage of expression.

To confirm cluster annotation, we assessed the expression of marker genes using the uniform manifold approximation and projection (UMAP)(**Fig. 2C**). After sub-clustering and doublet removal (**Methods**), we identified 3,625 vECs, our main cell-type of interest. These vECs constitute 5.4% of the total cells in our quality-controlled dataset. This proportion of vECs aligns with expectations for dermal vascular density, considering the removal of major epidermal populations (keratinocytes and basal cells) and using the endothelial cell enrichment method (**Methods**).

### Transcriptomic analyses of murin skin vECs

By performing differentially expressed gene analysis in vECs, we found a single gene that was differentially expressed after multiple testing correction (**Supplementary Table 1**). *Zbtb16* (zinc finger and TBT domain containing 16), also known as promyelocytic leukemia zinc finger (PLZF), was significantly overexpressed in vECs of the PA mouse group (**Fig. 3A**). In the other cell-types that we profiled by scRNAseq, we found no other genes that were significantly differentially expressed (**Supplementary Table 2**). While not the focus of our study give the limited sample size, we also performed sex-stratified differential gene expression analysis in vECs and found four genes (*Cyp1a1*, *Cfd*, *Car4*, *Gm11290*) down-regulated in female mice from the PA group (**Supplementary Fig. 1**). The role of these genes in response to PA or cellular stress in vECs is unknown, although Cyp1a1 expression is induced by shear stress (22) and Cfd has been implicated in several cardiovascular and metabolic diseases (23).

**Figure 3.**
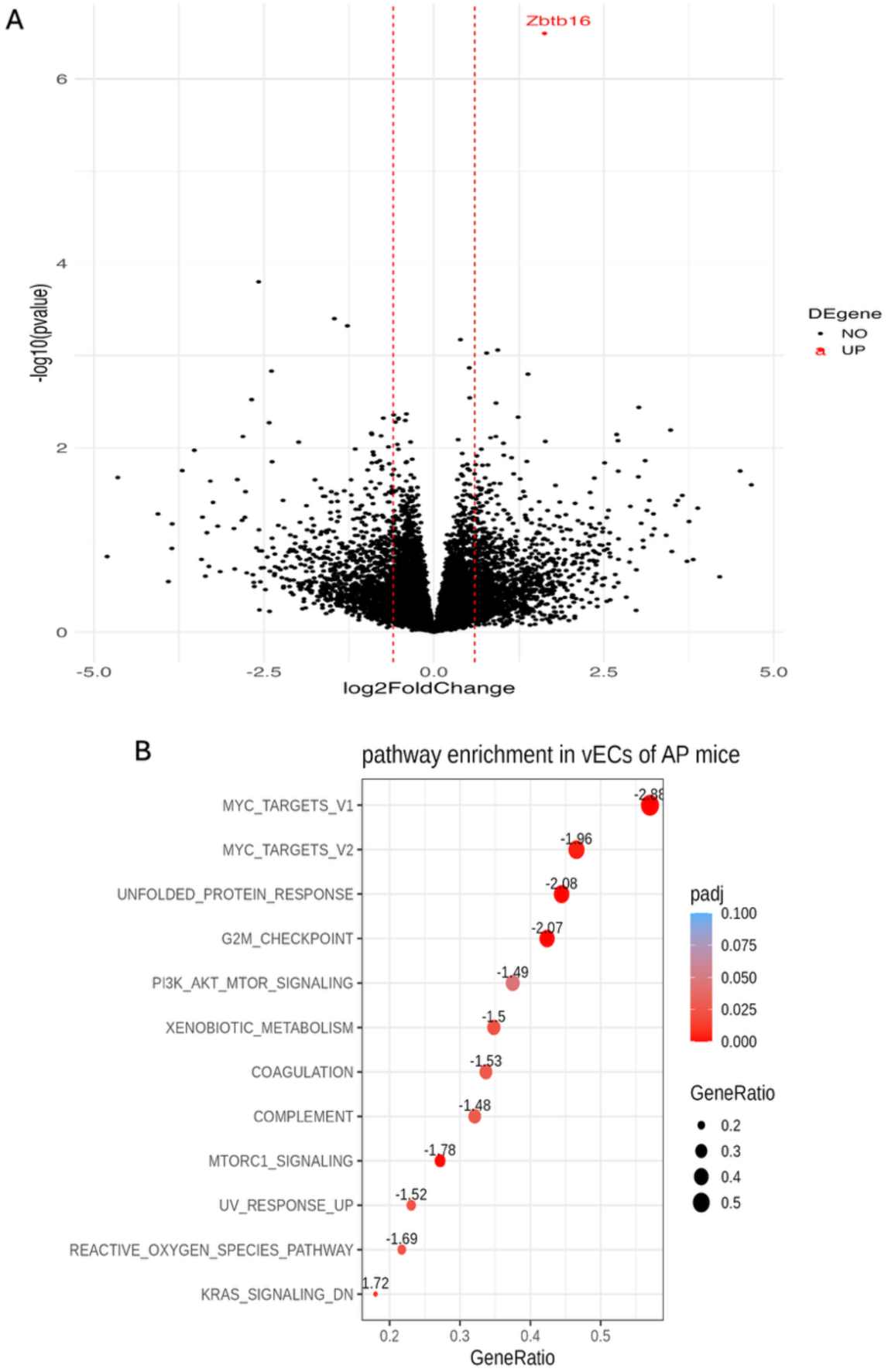
DiNerentially expressed gene and gene set enrichment analyses. (**A**) Volcano plot depicting di_erentially expressed genes. The analysis identified *Zbtb16* (zinc finger and BTB domain containing 16) as significantly overexpressed in vECs of the PA mouse group. (**B**) Gene Set Enrichment Analysis (GSEA) of vECs from murine skin comparing PA and sedentary groups. The analysis shows significant enrichments in pathways related to cell cycle regulation and stress response.

While we could only detect one gene that was significantly dysregulated in murine vECs upon exercise after correcting for multiple testing, we reasoned that several genes that reached nominal significance could nominate common biological pathways. GSEA highlighted enrichment of several key pathways related to cell cycle regulation, such as the “activation of KRAS-signaling-DN” pathway that captures genes that are downregulated when KRAS signalling is activated (**Supplementary Table 3**). GSEA results also revealed a consistent suppression of multiple pathways typically associated with cell proliferation (MYC Targets V1 and V2), stress response (Reactive Oxygen Species Pathway, Unfolded Protein Response, and UV Response Up), and metabolism (mTORC1 Signaling) in vECs of PA mice (**Fig. 3B**). The suppression of these pathways suggests a shift towards lower cellular stress in vECs of PA mice. Many genes implicated by the GSEA are correlated with the expression of the transcription factor *Zbtb16* (47 and 8 positively and negatively correlated, respectively, adjusted P-value <0.05) (**Supplementary Table 4**).

## DISCUSSION

In humans, regular PA increases nitric oxide production and bioavailability and reduces the production of pro-inflammatory cytokines and reactive oxygen species (24). These observations point towards an effect of PA on endothelial functions, but until recently it was challenging to distinguish the PA-triggered molecular changes that occur in vECs from those that occur in other cell-types of the blood vessels (e.g. smooth muscle cells, immune cells). In this study, we took advantage of scRNAseq to profile the transcriptome of vECs from the murine skin microvasculature in response to exercise.

Our analyses identified a single gene, *Zbtb16*, that is upregulated in skin vECs after exercise. *Zbtb16* encodes a poorly characterized transcription factor that has not been directly linked to PA response yet. Earlier work suggested that *Zbtb16* can suppress endothelial cell proliferation and angiogenesis (25,26), and that it is also involved in the endothelial response to various stimuli such as arsenic (27), far-infrared therapy (28), or pro-inflammatory TNFα treatment (29). Our analyses showed that *Zbtb16* expression is vECs is correlated with the expression of many genes implicated in biological pathways that respond to exercise (see below). Taken together, these results suggest that *Zbtb16* plays a critical role in vEC adaptation to cellular stress. Further experiments are now needed to dissect whether and how the *Zbtb16* transcriptional network contributes to improve endothelial functions in response to PA.

Acute exercise is generally associated with increased stress response in many tissues (30). In contrast, our GSEA suggest that a 1-month PA period leads to a suppression of stress-related pathways in vECs (**Fig. 3B**). The downregulation of stress response pathways, such as the Reactive Oxygen Species and Unfolded Protein Response pathways, indicates that PA may enhance vEC resistance to cellular stress. The dysregulation of genes implicated in the key mTOR pathway also suggests that the potential increased vEC resilience to stress after PA may be explained by more efficient cell metabolism (**Fig. 3B**). Consistently, *Zbtb16* has been implicated in the metabolic syndrome through an impact on mitochondrial functions, fatty acid oxidation and glycolysis (31,32).

Our study has two main limitations. First, we noted that a voluntary 1-month exercise exposure in young and healthy mice was not sufficient to improve vascular endothelial functions (**Fig. 1A-B**), although we observed expected changes in terms of weight loss (**Fig. 1C**) and plasma lipid profile (**Fig. 1D**). It is likely that a longer exercise treatment in mice could highlight other molecular mechanisms associated with the vascular benefits of PA. Second, despite our attempt to enrich for endothelial cells (**Methods**), vECs only represented 5.4% of all the cells that we profiled by scRNAseq, which is expected considering that the endothelium is a monolayer in blood vessels. This resulted in lower statistical power to identify genes for which PA has a modest impact on their expression levels. Because the skin is notoriously difficult to dissociate – an essential step in scRNAseq experiments – the development of more robust protocols to prepare skin sample will improve the yield of such experiments for rarer cell-types (33).

In conclusion, we profiled a transcriptomic dataset of the skin of voluntary exercising and sedentary mice, focusing on vECs. Our aim was to identify differentially expressed genes regulated by PA. We found that the transcription factor *Zbtb16* is highly upregulated in the skin vECs of PA mice and may participate in the adaptation of cellular stress. Because the skin is easily accessible, our study highlights the feasibility to extend this experimental protocol to better understand the vascular response of human subjects to PA.

## Supporting information

Supplementary Tables (Excel File with Multiple Sheets)

## DATA AVAILABILITY

The scRNAseq data discussed in this publication have been deposited in NCBI’s Gene Expression Omnibus and are accessible through GEO Series accession number GSE295877 (https://www.ncbi.nlm.nih.gov/geo/query/acc.cgi?acc=GSE295877).

## CODE AVAILABILITY

The code used to analyze the data and generate the figures is available upon request.

## ACKNOWLEDGEMENTS

This work was funded by the Montreal Heart Institute Foundation, the Joseph C. Edwards Foundation, the Canada Research Chair Program, and the Canadian Institutes of Health Research (Project #168902) to G.L..

## AUTHOR CONTRIBUTIONS

All authors conceived and designed the experiments. P.M. and P.H. collected the data and performed analyses. E.T. and G.L. secured funding and supervised the work. P.H., P.M., and G.L. wrote the manuscript with contributions from all authors.

## COMPETING INTERESTS

The authors declare that they have no competing interests.

## extended Figures

**Extended figure 1.**
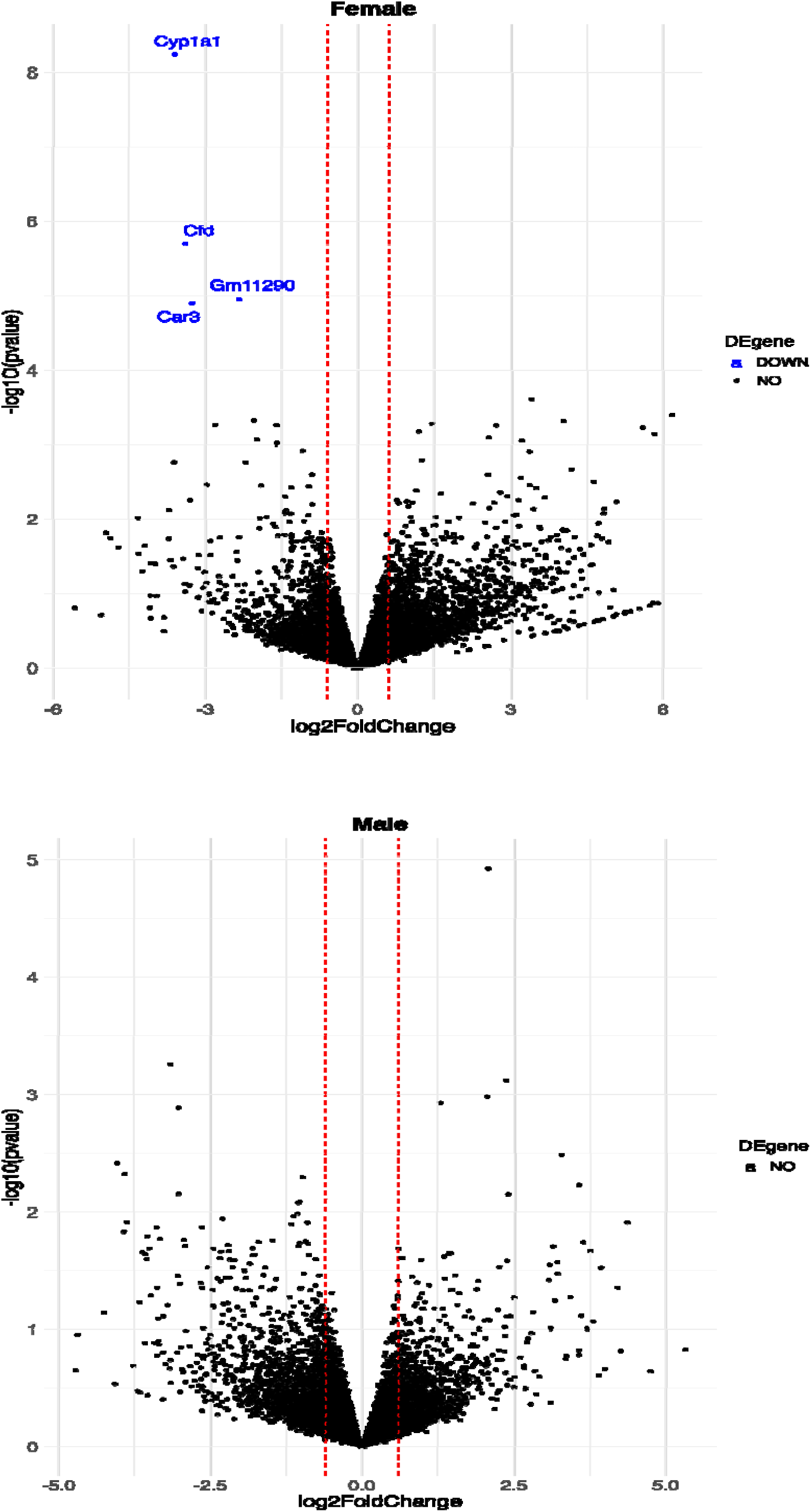
Sex-stratified differential gene expression analysis of vEC of PA versus SED mice.

## Notes

https://www.ncbi.nlm.nih.gov/geo/query/acc.cgi?acc=GSE295877

### Competing Interest Statement

The authors have declared no competing interest.

